# Cross-species proteomic and microRNA comparison of extracellular vesicles in human milk, cow’s milk, and infant formula products: moving towards next generation infant formula products

**DOI:** 10.1101/2023.02.23.529810

**Authors:** Natalie P. Turner, Pevindu Abeysinghe, Pawel Sadowski, Murray D. Mitchell

## Abstract

Milk and milk products such as infant formula (IF) play a fundamental role in serving the nutritional needs of the developing infant. Extracellular vesicles (EVs) in human (HM) and cow’s milk (CM) contain molecular cargo such as proteins and micro(mi)RNA that serve as functional messengers between cells and may be of importance to infant health. Here, we have developed a pipeline using advanced proteomics and transcriptomics to enable cross-species comparison of milk and IF EVs. EVs from HM, CM and IF were subjected to data-independent acquisition mass spectrometry and RNA-seq. Differentially abundant proteins (143) and miRNAs (514) were identified in HM and CM EVs, and CM EV proteins and miRNAs were preserved in IF EVs (∼20% protein; ∼90% miRNA). We foresee this work to be used in large scale studies to determine biologically relevant species-specific differences in milk EVs that could be leveraged to improve IF products.

## 1. Introduction

The first thousand days of life is a transient period of developmental plasticity unique to early life. Infant nutrition contributes significantly to establishing optimal health during this time and influences the overall risk of adult diseases such as asthma, diabetes and hypertension (Davies et al., 2016). Major components of human milk (HM) include proteins such as lactoferrin, immune factors, immunoglobulins, cytokines, and growth factors essential for growth and development in the newborn (Ballard & Morrow, 2013). The breastfed infant is also exposed to microbes in HM that assist in airway and gut microbiota population (Le Doare et al., 2018). Intense interest in the composition of HM has been spurred, in part, by a growing demand for infant formula (IF) products. While most IF products have a protein base derived from cow’s milk (CM), the similarities and differences to HM are not well-understood. To align the nutritional advantages of HM with IF, distinguishing differences between the macronutrients and *<u>micronutrients</u>* in various milk types is required to move towards next generation IF recipes.

Mass spectrometry-based proteomics has been used for the analysis of milk proteomes from different animal species, such as cow, goat, camel, yak, and HM in several studies (Manoni et al., 2020; Yao et al., 2021). Proteins found in abundance in HM over other species confer significant health benefits, including increased bacterial resistance (Yao et al., 2021), reduced incidence of gastroenteritis (Stevens et al., 2000), and protection against rotavirus infection (Newburg et al., 1998). Remarkably, there are oligosaccharides found only in HM that contribute to a diversified microbiome and as such the breastfed infant gastrointestinal tract is host to vastly different microflora than formula-fed infants (Ballard & Morrow, 2013; Le Doare et al., 2018). The establishment of gut microbiota is therefore crucial to gastrointestinal health in early life, a process which may be altered in formula-fed infants.

Separate to macronutrients, milk micronutrients also play an important and yet understudied role in infant health. Among the micronutrients in milk exists a class of lipid membrane-bound nanoparticles that contribute to the nutritional uptake of milk components in the gut; extracellular vesicles (EVs). EVs are abundant in milk and contain bioactive molecular cargo including, but not limited to, proteins, nucleic acids and microRNAs (miRNAs) (Abels & Breakefield, 2016). They are released by cells into the local and distant milieu and taken up by recipient cells, delivering their contents and effecting changes in the cell (Jiang et al., 2021). Three major classes of EVs are ectosomes/microvesicles (100 – 1000 nm), which are shed directly from the plasma membrane, exosomes (50 – 150 nm), originating from the endocytic pathway, and apoptotic bodies (1 – 5 µm), released by cells undergoing cell death (Jeppesen et al., 2019). While EVs contribute to the total milk proteome and transcriptome, to effectively study their cargo they must be enriched from milk by depleting other abundant milk components such as milk fat globules (MFG) and casein (K. Vaswani et al., 2019).

Milk EVs can survive the harsh conditions of the upper digestive tract to pass into the intestines, where they can be taken up by the intestinal epithelium (Samuel et al., 2021). EVs deliver their cargo directly to the intestinal epithelia and are not metabolised beforehand; thus, extracting proteins from intact EVs is fundamental to understanding their function. The effects of HM and CM EVs on the gut in various *in vivo* and *in vitro* models suggest that exposure to gut cells suppresses inflammation while promoting positive changes in gut cell morphology, microbiota, and increasing nutrient uptake (Du et al., 2021; Tong et al., 2021). Specifically, the milk proteins butyrophilin subfamily 1 member A1 (BTN1A1), lactadherin (LDH), xanthine dehygrogenase/oxidase (XDH) and perilipin-2 (PLIN2) are now known to be EV-associated (Samuel et al., 2017). These immunomodulatory proteins are common to HM and CM EVs, however the differential abundance of these proteins in milk between species is currently unknown and their specific roles when EV-associated are unclear. Regarding the role of miRNA in the gut, intraperitoneal administration of HM EV-derived miR-148a-3p was found to reduce inflammation and improve the gut epithelial tight junctions in a mouse model of necrotising enterocolitis (Guo et al., 2022). However, reduced levels of EV miR-148a have been identified in commercially available IF products compared to HM (Golan-Gerstl et al., 2017).

Despite the obvious importance of EVs to milk composition, limited work has been performed to characterise the qualitative and/or quantitative differences between the EV miRNA and proteins found in HM, CM and IF. To our knowledge there have been no proteomic studies to date on EVs derived from IF products. Understanding the differences in proteomic and miRNA content in milk EVs across species could be leveraged to generate IF products optimised according to HM EVs, therefore reducing the nutritional gap between breastfed and formula-fed infants.

Considering the positive effects milk EVs exert on the gut microbiota and their ability to improve nutrient uptake, we aimed to develop a workflow that would permit comparison of the proteomic and transcriptomic profiles of EVs derived from HM, CM and IF. First, we performed proteomic and transcriptomic analysis on small EVs enriched from HM and CM EVs via data-independent acquisition (DIA) mass spectrometry and next-generation miRNA sequencing. Utilising bioinformatics tools, we were able to generate cross-species libraries used for quantitative and qualitative data analysis and quantify differences in shared proteins and miRNAs between species. As a means of understanding the potential pitfalls of IF processing on EVs contained within IF products, we report for the first time the successful recovery of intact EVs from 9 different lyophilised CM-based IF products, which were subsequently subjected to EV characterisation, proteomic and miRNA analysis for comparison with HM and CM EVs.

## 2. Materials and Methods

### 2.1 Ethics approvals

For human milk samples, donors gave informed consent and ethics were approved by the QUT Office of Research Ethics & Integrity (OREI; approval # 1900000892). Cattle milk samples were collected according to ARRIVE guidelines (QUT OREI, animal ethics approval # 1800000485, QV ref # 83812).

### 2.2 Sample Collection

Human milk was self-collected using an electric breast pump by two healthy mothers with normal pregnancies and uncomplicated term deliveries (38 – 40 weeks gestation). Milk was collected between 4 – 6 months post-partum, transferred into storage bags, sealed and stored at -20 °C. Frozen milk samples were transferred to -80 °C within 4 weeks of collection.

Unpasteurized cow’s milk was collected at a local dairy farm in Gatton, QLD, from Holstein-Friesian (*Bos taurus*) dairy cows (*n* = 2) as previously described (K. Vaswani et al., 2017). Milk was aliquoted (50 mL/aliquot) and stored at -80 °C.

Lyophilised infant formula products were obtained from commercial manufacturers Nutricia (Aptamil), Mead Johnson (Enfamil/Enfinitas) and Nestle (NAN). Each formula range consisted of a 0– 6-month (0-6), 6-12-month (6-12) and 1 year+ (1Y) product, which were reconstituted per manufacturer’s instructions on the same day EV isolation commenced.

### 2.3 Experimental Design

To compare the milk proteomes between species and reduce differences based on individual biological variability, HM (*n* = 2) and CM (*n* = 2) samples were pooled separately to total volume 180 mL (HM) and 200 mL (CM), based on the availability of each milk type. IF products were treated as biological replicates based on the brand of IF, which created 3 groups of IF containing 3 biological replicates each (see Table 1).

**Table 1:**
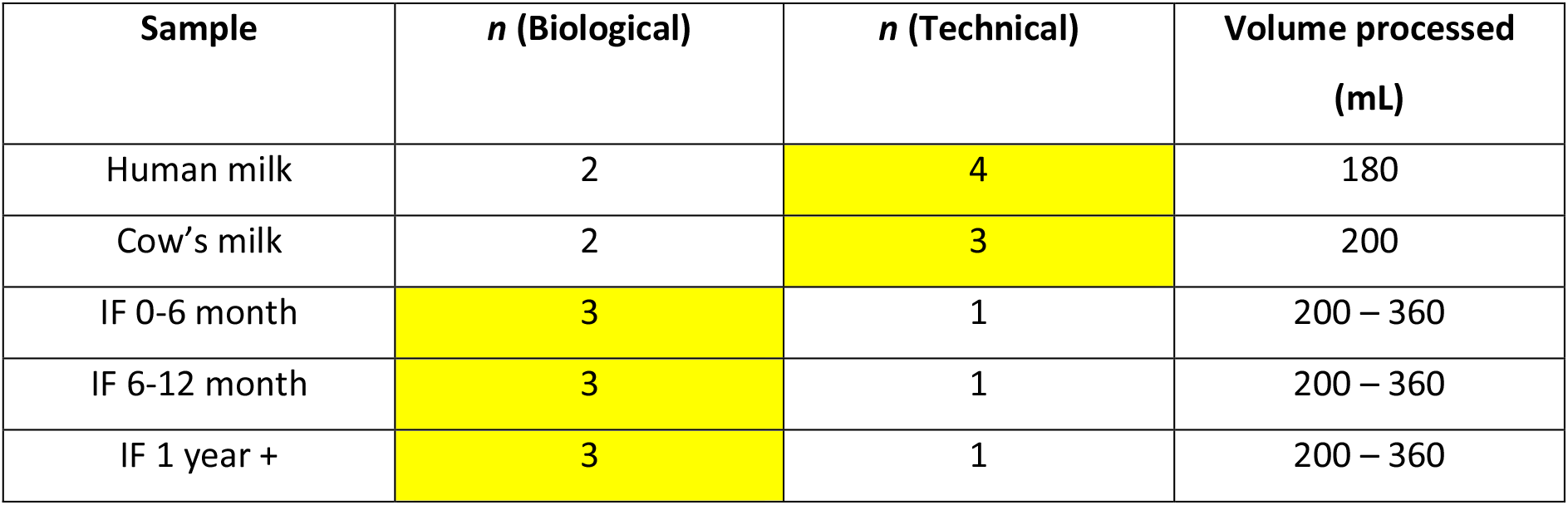
Sample information for milk and infant formula (IF) products used in this study. IF volumes were adjusted based upon the recommendation from the manufacturer, where volume processed = recommended volume for 1 bottle feed. Yellow areas indicate *n* values used for statistical analysis.

| Sample | *n* (Biological) | *n* (Technical) | Volume processed (mL) |
| --- | --- | --- | --- |
| Human milk | 2 | 4 | 180 |
| Cow's milk | 2 | 3 | 200 |
| IF 0-6 month | 3 | 1 | 200 – 360 |
| IF 6-12 month | 3 | 1 | 200 – 360 |
| IF 1 year + | 3 | 1 | 200 – 360 |

### 2.4 Extracellular vesicle isolation from milk and infant formula products by Ultracentrifugation (UC) and Size-exclusion chromatography (SEC)

#### 2.4.1 UC

EV isolation for HM, CM and IF was performed as previously described (K. M. Vaswani et al., 2021). Briefly, human and bovine milk samples were thawed on ice the same day EV isolation commenced. Human milk samples were pooled, mixed thoroughly, and separated into 50 mL aliquots for processing. A total of 200 mL of milk/species was processed for EV isolation. For IF samples, following the manufacturer’s instructions for formula weight equivalent to the recommendation for one bottle feed, samples were reconstituted with sterile de-ionised H2O (ranging from 200 mL – 360 mL) and mixed thoroughly, before being aliquoted into 50 mL falcon tubes for EV isolation. Samples were centrifuged at 3,000 × *g* at 4 °C to remove cells, cell debris and particulate matter. Supernatants were aspirated and transferred to new falcon tubes, with care being taken not to disturb the pellets. To precipitate casein, supernatants were mixed 1:1 with 0.25M EDTA and incubated for 15 min on ice, then centrifuged at 12,000 × *g*, 35,000 × *g* and 70,000 × *g* for 60 min each at 4 °C to remove MFGs, casein, apoptotic bodies and microvesicles. After the final centrifugation step, supernatants were filtered through a 0.22 µm filter (Corning Co-star, Mulgrave, Victoria), before being transferred into 32.4 mL Optiseal Polypropylene Tubes (Beckman Coulter, Brea, CA, USA). Samples were centrifuged at 100,000 × *g* for 2 hr at 4 °C (Type 50.2 Ti Fixed angle ultracentrifuge rotor, Beckman Coulter, Brea, CA, USA). The supernatants were aspirated, and pellets resuspended in 500 µL Dulbecco’s phosphate buffered saline, pH 7.0 – 7.3 (DPBS; Gibco, Thermofisher Scientific, Brisbane, Australia). Pellets from each sample type were combined, vortexed, and stored overnight at 4 °C prior to SEC.

#### 2.4.2 SEC

The resuspended pellets obtained from UC were enriched for small EVs (diameter <200 nm) by loading onto qEV original 70 nm size-exclusion columns (Izon Science Ltd, New Zealand) in 500 µL aliquots as per manufacturer’s instructions. A total of sixteen 500 µL fractions were collected per sample as previously described (K. Vaswani et al., 2017) and fractions 7 – 10 pooled as the EV-containing fractions. This resulted in four replicates of HM EVs, three replicates of CM EVs, and three replicates of each IF brand and age group (Table 1). Pooled EV fractions were stored at -80 °C until required for characterisation experiments and downstream analyses.

### 2.5 EV Characterisation

#### 2.5.1 Western Blot

CM, HM and IF pooled EV fractions were assayed for total protein according to manufacturer’s instructions (Micro BCA protein assay kit, Thermofisher Scientific, Brisbane, Australia) and a volume equivalent to 20 µg total protein was aliquoted for western blot (WB) analysis. Human JEG-3 choriocarcinoma placental cell lysate (10 µg) was used as a positive or negative control as appropriate (Walton et al., 2013). Samples were resolved by protein electrophoresis as previously described (N. P. Turner et al., 2022). Briefly, all samples were placed in a vacuum centrifuge (SpeedVac, Eppendorf, Sydney, Australia) until dry. A master mix containing 4x LDS sample buffer, 10x reducing agent and de-ionised H2O was made to final concentration 1x (NuPAGE, Thermofisher Scientific, Brisbane, Australia). Thirty µL of master mix was added to each dried sample, vortexed and centrifuged. Samples were sonicated for 2 min then transferred to a heating block and reduced at 70 °C for 10 min. Reduced samples were transferred to a 4 – 12% Bis-tris mini protein gel (Thermofisher Scientific, Brisbane, Australia) and proteins resolved by electrophoresis. The transfer of protein bands to a polyvinylidene fluoride membrane was performed using the Transblot Turbo system (Bio-Rad Laboratories Pty Ltd., Sydney, Australia), as previously described. Membranes were blocked in Intercept (PBS) blocking buffer (Li-COR, Mulgrave, Australia) for 1 hr at RT with gentle rocking. After blocking, membranes were transferred to primary antibody diluted in 1:1 phosphate buffered saline (PBS): blocking buffer, with Tween-20 added to final concentration of 0.1% (PBST) and incubated overnight at 4 °C; anti-Bovine serum albumin (ALB) (1:5000, Rabbit polyclonal (Abcam, Melbourne, Australia); recombinant anti-Flotillin-1 (FLOT-1) (1:1000, Rabbit monoclonal (Abcam, Melbourne, Australia); recombinant anti-Tumour susceptibility gene 101 (TSG101) (1:1000, Rabbit monoclonal (Abcam, Melbourne, Australia); anti-CD9 (1:1000, Mouse monoclonal (Novus Biologicals (Invitro technologies), Melbourne, Australia); anti-CD81 (1:1000, Rabbit polyclonal (Novus Biologicals (Invitro technologies), Melbourne, Australia); anti-Glyceraldehyde-3-phosphate dehydrogenase (GAPDH) (1:5000, Rabbit polyclonal (Novus Biologicals (Invitro technologies), Melbourne, Australia); anti-Calnexin (CALNX) (Rabbit polyclonal (1:2000, Rabbit polyclonal (Novus Biologicals (Invitro technologies), Melbourne, Australia); anti-Syntenin 1 (SYN-1) (1:1000, Rabbit polyclonal (Novus Biologicals (Invitro technologies), Melbourne, Australia). All primary antibodies were listed as reactive in human and cow by the manufacturer. The next day, membranes were transferred to clean membrane trays and washed 4x in PBST. Washed membranes were incubated with secondary antibody in the dark for 1 hr at RT with gentle rocking (Goat anti-Rabbit IgG, 1:20,000, Li-COR, Mulgrave, Australia; Goat anti-Mouse IgG (H+L), 1:20,000, Cell signalling Technology, Melbourne, Australia). Membranes were washed 4x with PBST, 1x with PBS and imaged on a fluorescent scanner (Li-COR, Mulgrave, Australia) at 700 and 800 nm. Images were processed using Image Studio Lite v5.2.

#### 2.5.2 Nanoparticle Tracking Analysis

Particle size and concentration measurements were performed on a NanoSight NS300 instrument (Malvern Panalytical, Sydney, Australia). Instrument calibration was performed using 100 nm polystyrene beads (Malvern Panalytical, Sydney, Australia). EV pooled samples were diluted in filtered DPBS until a concentration of 30 – 70 particles per frame was reached. Camera level was set to 13, detection threshold set to 5, and three captures of 30 sec each were performed at a flowrate of 300 µL min^-1^. Processed results were entered into a .xlsx file and exported to GraphPad Prism (v9.5.0) (GraphPad, San Diego, CA, USA) for graphing and statistical analysis as previously described (N. P. Turner et al., 2022).

#### 2.5.3 Transmission Electron Microscopy

Pooled EV samples were visualised by transmission electron microscopy (TEM), performed by staff at the Central Analytical Research Facility microscopy lab (Queensland University of Technology, Brisbane, Australia). EV samples were diluted 1:1 with de-ionized H2O, mounted onto formvar coated 200 Cu mesh grids and imaged as previously described (N. P. Turner et al., 2022).

### 2.6 Proteomic Sample Preparation and Analysis

#### 2.6.1 Protein Digestion

A volume corresponding to 50 µg of total protein was used for proteomic sample preparation for each sample type. EV proteins were extracted using 1% (HM and CM) or 2% (IF) w/v sodium deoxycholate lysis buffer and processed as previously described (N. Turner et al., 2022). In brief, following the addition of lysis buffer, samples were sonicated for 2 min in an ice bath and incubated on ice for 20 min with gentle rocking. Samples containing lysis buffer were loaded onto 30K omega membrane centrifugal filters (PALL, Brisbane, Australia) and centrifuged. All centrifugation steps were performed at 14,000 × *g* for 15 min at 21 °C. On-filter reduction was performed with 200 µL of 25 mM DTT for 1 h at RT with gentle agitation. Filters were centrifuged and membranes washed once with 200 µL of 8M urea. Alkylation was performed by adding 100 µL of 50 mM iodoacetamide to the filters and incubating in the dark for 20 min at RT with gentle agitation. The filters were centrifuged and washed twice with 200 µL of 8M urea, followed by two additional washes with 200 µL of 100 mM ammonium bicarbonate pH 8.0 (AMBIC). Modified trypsin (Promega, Madison, WI, USA) stock was diluted in 100 mM AMBIC (0.1 µg/µL) and added to the filters at an enzyme:protein ratio of 1:50. Overnight digestion was performed at 37 °C in a humidified chamber. The next day, filters were transferred to clean 1.5 mL protein lo-bind tubes (Eppendorf, Sydney, Australia) and centrifuged to collect peptides. One more elution was performed by adding 60 µL AMBIC to the filters and centrifuging as per previous step.

#### 2.6.2 Desalting

Peptides were acidified by adding an appropriate volume of 4% trifluoroacetic acid (TFA) to a minimum final concentration of 0.1%. HM and CM peptides resulting from the previous step were desalted using peptide desalting spin columns (Thermofisher Scientific, Brisbane, Australia), according to manufacturer’s instructions. IF peptides were desalted using double strong cation exchange (SCX) Stagetips as previously described (N. Turner et al., 2022). Desalted peptides were dried in a vacuum concentrator (Concentrator plus, Eppendorf, Sydney, Australia) and resuspended in 16 µL indexed retention time (iRT) buffer (Biognosys-11).

#### 2.6.3 Peptide Assay

Peptide samples were assayed for peptide concentration with Pierce™ Quantitative Colorimetric Peptide Assay (Thermofisher Scientific, Brisbane, Australia), according to the manufacturer’s instructions. Samples were normalised for peptide concentration before submission for MS analysis by adding an appropriate volume of iRT buffer.

#### 2.6.4 Liquid Chromatography-Mass Spectrometry

Four µg of each peptide sample was analysed on TripleTOF 5600+ QqTOF instrument (SCIEX) configured for microflow applications as previously described (N. Turner et al., 2022). Chromatographic separation involved a 68 min linear gradient of 3-25% mobile phase B followed by 5 min linear gradient of 25-35% mobile phase B. After peptide elution, the column was flushed with 80% mobile phase B for 3 min and re-equilibrated with 97% A for 8 min before the next injection. For spectral library generation (qualitative runs), a non-fractionated pooled biological quality control (PBQC) sample was analysed using a Data-Dependent Acquisition (DDA) approach. For quantitation purposes, individual samples were analysed using a variant of Data-Independent Acquisition (DIA) approach termed <u>S</u>equential <u>W</u>indow <u>A</u>cquisition of all <u>Th</u>eoretical <u>M</u>ass <u>S</u>pectra (SWATH-MS). Additionally, a minimum of two replicate injections of PBQC sample were acquired using the SWATH method during analysis of the quantitative batch for quality control purposes.

##### 2.6.4.1 Data-dependent Acquisition (DDA)

The PBQC and individual peptide samples were subjected to TOF MS scans collected at a resolution of 30,000 over a *m/z* range of 400 -1,250 for 0.25 s. This was followed by high sensitivity TOF MS/MS scans over a *m/z* range of 100 - 1,800 on up to 30 of the most abundant peptide ions (0.05 s per each scan) that had intensity greater than 150 counts per second (cps) and a charge state of 2-5. The dynamic exclusion duration was set at 15 s. All other parameters (ion fragmentation, collision energy, declustering potential (DP)) were as previously described (N. Turner et al., 2022). Information gathered from these TOF MS scans were used to determine the species-specific *m/z* range for TOF MS scans used in the subsequent SWATH-MS workflow.

##### 2.6.4.2 Data-independent Acquisition (SWATH-MS)

All peptide samples were analysed by SWATH-MS. SWATH-MS was based on the approach published by Gillet *et al*. (2012) with modifications (Gillet et al., 2012). A high-resolution (30,000) TOF MS scan was collected over a range of 400 - 900 *m/z* (*Bos taurus*) or 400 – 1000 *m/z* (*Homo sapiens*) for 250 ms and from that range precursors were selected for subsequent high-sensitivity TOF MS/MS scans using 60 custom variable Q1 windows. The mass range of each product ion scan was 100 – 1800 and the dwell time was set at 50 ms and 30 ms, respectively. The collision energy for each window was set using the collision energy of a 2+ ion centred in the middle of the window (equation: 0.0625 × *m/z* -3 in the case of TripleTOF 5600+ with a spread of 5 eV. The DP was set to 80 V and the remaining gas and source parameters were adjusted as required. The cycle time for *Homo sapiens* and *Bos taurus* was 2.2 s and 3.1 s, respectively, which resulted in a minimum of 6 data points per each peptide ion peak. Full details of the SWATH variable windows used for each species can be found in Supplementary file 1.

##### 2.6.4.3 In silico Peptide Spectra Library Generation

Common contaminants (cRAP; http://ftp.thegpm.org/fasta/cRAP/) and iRT peptide sequences were appended to reference proteome sequences in FASTA format for *Bos taurus* and *Homo sapiens* downloaded as reference proteomes from Uniprot (23,847 sequences and 81,837 sequences, respectively. Downloaded December 21^st^, 2021). These proteome sequences were provided to DIA-NN software (Demichev et al., 2020) either individually (species-specific libraries) or combined (cross-species library) were for *in silico* digestion and peptide spectra prediction. The parameters were set as follows: Deep learning enabled; FDR 0.01; min peptide length = 6; max peptide length = 30; precursor min *m/z* = 400; precursor max *m/z* = 1000; min fragment *m/z* 100; max fragment *m/z* 1800; Protease = Trypsin/P; Missed cleavages = 1; Max number variable modifications = 3; N-terminal methionine excision enabled; cysteine carbamidomethylation set as fixed modification; Methionine oxidation enabled as variable modification.

##### 2.6.4.4 SWATH Data analysis

Raw data files containing SWATH-MS scans were analysed using DIA-NN software against in silico libraries with additional settings as follows: (MS2) mass accuracy = 15 ppm; MS1 mass accuracy = 20 ppm; remove likely interferences enabled; use isotopologues enabled; Neural network classifier = Double-pass mode; Protein inference = Genes; Quantification strategy = Robust LC (high accuracy); Cross-run normalisation = RT and signal-dependent (experimental); Library generation = Smart profiling. The generated individual species report files were used for protein identification, whereas cross-species report files (.tsv) were imported into R Studio for peptide quantitation using packages MsStats, diann and dplyr for Volcano plot generation (data normalisation = equalise medians; significance = adjusted *p* < 0.05; fold change cut-off = 2). R packages ggbiplot and tidyverse were used to conduct principal component analysis (PCA). For cross-species protein quantitation, filtering was applied in R to include proteins common to both species based on peptide homology, and differential analysis was performed using homologous sequences only. For protein identification, protein groups identified at 0.01 FDR were extracted directly from DIA-NN output and filtered to include proteins quantified in all replicates.

##### 2.6.4.5 Gene ontology

Proteins identified by MS were subjected to gene ontology (GO) analysis using PANTHER (http://www.pantherdb.org/) as previously described (N. Turner et al., 2022). For cross-species comparison, differentially abundant proteins common to HM and CM were searched against organism *Homo sapiens* and inspected manually for differences in cellular component, biological pathway, molecular function, and biological process. Data files containing GO information were downloaded from PANTHER as .txt files and imported into Excel. For CM and IF comparison, proteins identified in both sample types were searched against organism *Bos taurus* and assessed for differences in the same way.

#### 2.6.5 MicroRNA Sample Preparation and analysis

##### 2.6.5.1 MiRNA isolation and purification

A volume of EV sample equivalent to 50 µg of total protein was used for EV miRNA isolations. EV samples were processed for miRNA isolation using TRIzol (ThermoFisher Scientific, Brisbane, Australia), Chloroform (Sigma-Aldrich (Merck), Melbourne, Australia) and miRNeasy mini kit (QIAGEN, Melbourne, Australia) as previously described (Abeysinghe et al., 2021). Briefly, EV samples were incubated with TRIzol at room temperature (RT) for 5 minutes (2.5 volume of sample: 7.5 volume of TRIzol). Next, Chloroform was added (2.5 volume of sample: 1.5 volume of Chloroform), mixed thoroughly for 15 s and incubated at RT for 2 min. After a phase separation was visible, the sample was centrifuged for 15 min at 4°C, at 12,000 × *g*. The upper aqueous layer was carefully transferred to a new microcentrifuge tube and 1.5 volumes of 100% ethanol was added. This mixture was passed through a miRNeasy mini column and miRNA was isolated according to the manufacturer’s protocol. RNase free water supplied by the miRNeasy mini kit was passed through the column for twice consecutively (each 40 µL) to collect 80 µL of EV miRNA sample. Enriched sEV miRNA samples were shipped in dry ice to Australian Genome Research Facility (AGRF; Brisbane, Australia) for next generation miRNA sequencing immediately after sEV miRNA isolations.

##### 2.6.5.2 MiRNA Next-Generation Sequencing

Sequencing was performed by the AGRF as previously described (Abeysinghe et al., 2021). Briefly, miRNA library preperation was conducted by NEBNext® Multiplex Small RNA Library Prep Set for Illumina® and next, Illumina Novaseq S1 next-generation sequencing platform was utilized for single end 100bp sequencing. Image analysis was performed in real time by the NovaSeq Control Software (NCS) v1.7.5 and Real Time Analysis (RTA) v3.4.4, running on the instrument computer. Then the Illumina bcl2fastq2.20.0.422 pipeline was used to generate the untrimmed 100bp sequence data in FastQ format. The quality and integrity of FastQ sequence files were checked using checksum information generated by Exactfile (http://www.exactfile.com/).

##### 2.6.5.3 Analysis of miRNA sequence data

The quality above Q30 >78.19% bases were screened using FastQC and next, the reads were trimmed to remove overrepresented sequences, and length filtered using Trim_Galore! to be between 14 and 38 base pairs long for the Illumina adapter (AGATCGGAAGAGCACACGTCTGAACTCCAGTCAC). The cleaned sequence reads were then aligned against the *Bos taurus* genome (Build version UMD3.1) for CM and IF. *Homo sapiens* (Build version GRCh38.p14) was used to align reads for HM. The STAR aligner (v2.5.3a) was used to map reads to the genomic sequences (https://github.com/alexdobin/STAR/blob/master/doc/STARmanual.pdf) and alignment files were in BAM format. The counts of reads mapping to each known miRNA were identified using unitas (https://sourceforge.net/projects/unitas/). The differential gene expression was performed using edgeR (version 3.30.3) of R package 4.0.3 (https://bioconductor.org/packages/release/bioc/html/edgeR.html). Trimmed means of M values (TMM) method of edgeR was used to normalize the counts between the samples.

False discovery rate (FDR) analysis was performed to adjust the p-value for multiple hypothesis testing (Benjamini-Hochberg adjustment). FDR value of less than 0.05 (FDR < 0.05) was selected as the threshold to select statistically significant DE miRNA expressed between the groups for further analysis.

##### 2.6.5.4 miRNA target prediction and functional enrichment analysis of miRNA targets

The online target prediction tools miRwalk (http://mirwalk.umm.uni-heidelberg.de/) and miRnet https://www.mirnet.ca/) were used for qualitative analysis of differentially expressed miRNA resulting from quantitative analysis. To focus on the effects of EV miRNA on the human infant gut, miRNA target predictions were performed using *Homo sapiens* as the species.

## 3. Results and Discussion

### 3.1 Particle Size and Concentration

To confirm the recovery of particles in the small EV (< 200 nm) size range from HM and CM, we compared mean particle sizes of pooled EV fractions between these two groups. Particle size did not vary significantly between HM and CM EVs, with mean particle diameter ranging from ∼100 – 125 nm (Supplementary file 2, Figure S1A). Particle concentration was higher in CM than HM, which matches results previously reported by our group using UC+SEC as the method of EV isolation (Supplementary file 2, Figure S1B) (K. Vaswani et al., 2019). Moreover, particles obtained from IF samples were similar in their size (mean ∼80-100 nm) and particle concentration (×10^10-11^ particles/mL) to HM and CM. Total particle and protein yields and calculated particle/µg protein ratios are given in Table S1 (Supplementary file 2). The particle distribution plots obtained by NTA indicate that there is a high degree of similarity between HM and CM EVs, while IF EVs show greater variability and increased abundance of smaller particles ∼30 nm in size (Supplementary file 3).

TEM images of milk samples show vesicles of various sizes in the classical cup shape, with most distinguishable intact particles ranging from ∼30 nm – 200 nm (Supplementary file 4, Figure S1). This is consistent with previous findings in which we established that the combination of EV isolation methods used in this study results in a heterogenous mixture of particles < 200 nm diameter (K. Vaswani et al., 2019). IF samples also contained distinct cup-shaped particles of ∼50 – 200 nm in size and appeared to have more debris or non-vesicular material than HM and CM (Supplementary file 4, Figure S2). This is supported by the NTA data whereby protein aggregates are detected as smaller particles ≤30 and may indicate non-EV particle carry over from the IF production process. As this is the first study to recover EVs from IF products, future studies may assess the effects of different EV enrichment strategies on sample purity if this is of importance to the study design. As the CM EV proteome is well-established, comparison of the IF EV to the CM EV proteome can be achieved in this case without requiring an IF EV sample of high purity.

### 3.2 Enrichment of EV Protein Markers in HM, CM and IF

We first established the abundance of EV-enriched protein markers in milk and IF EVs to characterise the populations of EVs obtained. The exosomal proteins FLOT-1, TSG101 and CD81 were detected in HM, CM and IF EVs by WB and MS (Figure 1 and Supplementary file 5). CD9 was detected in CM and IF EVs by WB and MS, and in HM EVs by MS in this and other studies (van Herwijnen et al., 2016). Interestingly, CD9 abundance was quite consistent across all IF EV samples, although slightly reduced compared to CM, which suggests CD9-positive EVs are well-preserved in IF products. ALB was detected in IF but not in CM EVs by WB, therefore the presence of ALB in IF is likely carryover from the IF production process. SYN-1 is a putative universal exosome marker (Kugeratski et al., 2021) and was detected by MS in all samples, with additional validation by WB in HM and CM EVs (Supplementary file 5). FLOT-1 abundance by WB was similar between HM and CM EVs, however there was marked FLOT-1 signal reduction in IF EVs across all samples. FLOT-1 is a lipid raft protein that incorporates into the inner EV membrane during endosomal sorting and is considered to be a part of the inner leaflet or core EV proteome (Cvjetkovic et al., 2016); as such, changes to FLOT-1 abundance in IF should be interpreted with caution, as it may be indicative of a reduction of FLOT-1-positive EVs following IF processing rather than alterations to the protein itself.

**Figure 1:**
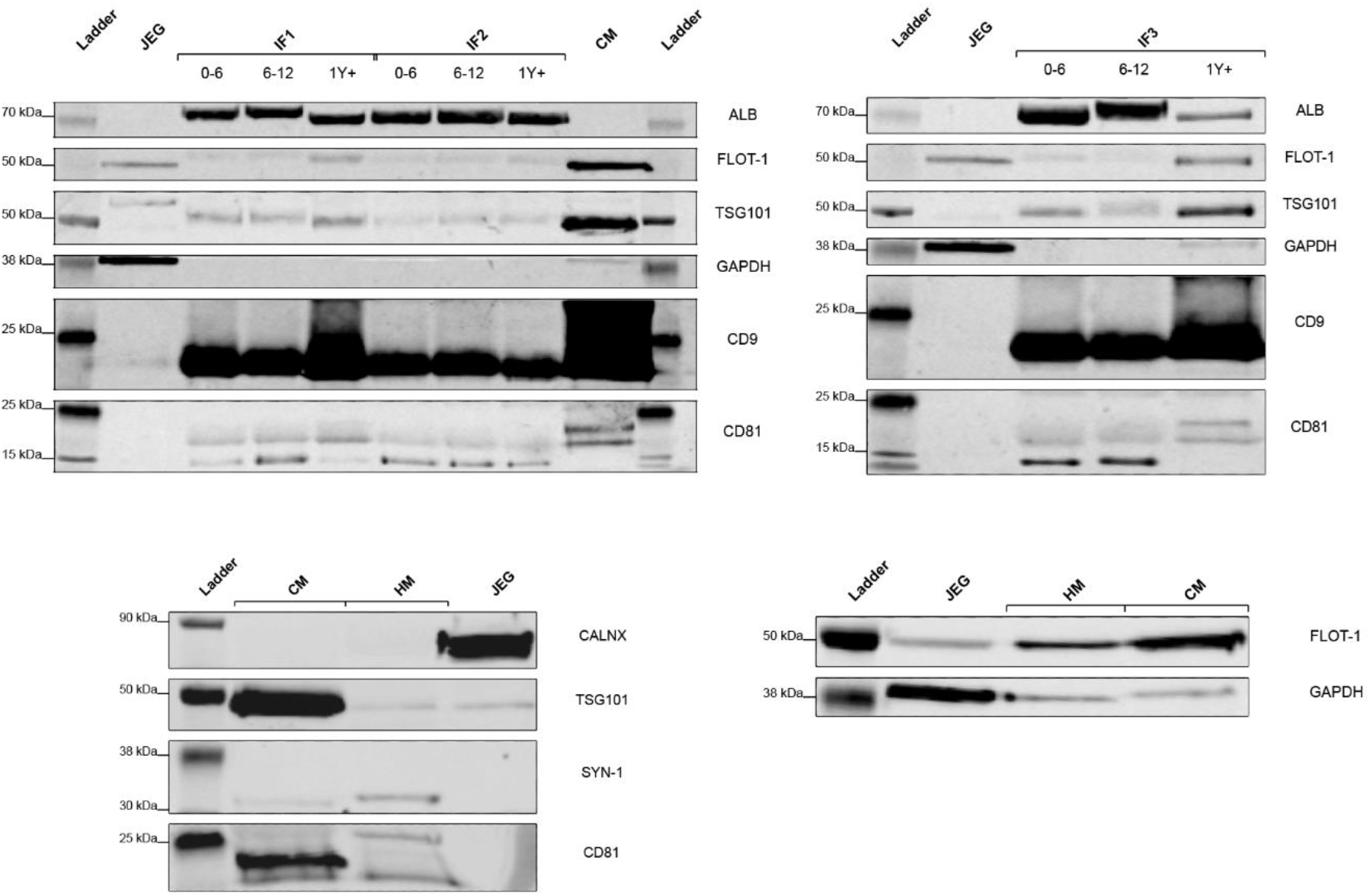
Western blot analysis of extracellular vesicle samples enriched from human milk (HM), cow’s milk (CM), and infant formula (IF) products. Top panels: Pooled EV fractions 7 – 10 enriched from infant formula (IF); 0–6-month (0-6), 6–12-month (6-12), and 1 year plus (1Y+) preparations; Pooled EV fractions 7 – 10 enriched from CM. Bottom panels: Pooled EV fractions enriched from HM and CM. JEG = JEG-3 cell lysate (+/-control); IF1 – 3 = Infant formulas from three different manufacturers; ALB = albumin; CALNX = Calnexin; GAPDH = glyceraldehyde-3-phosphate dehydrogenase; FLOT-1 = flotillin-1; SYN-1 = syntenin-1; TSG101 = tumour susceptibility gene 101.

The endoplasmic reticulum-associated protein CALNX is commonly used as a marker of cell protein contamination in EV preparations and was not detected in HM and CM EV samples, however cytosolic marker GAPDH was detected by WB and MS (Supplementary file 5). *Gapdh* has been reported as a housekeeping gene constitutively expressed in small EVs, while the GAPDH protein has recently been described as a major driver of EV biogenesis and promoter of EV clustering via surface binding to the EV membrane (Dar et al., 2021). GAPDH was not detected in 8 out of 9 IF samples by WB (Figure 1), however it was detected at low levels by MS. It is feasible that IF processing may disrupt and/or denature externally bound proteins, including GAPDH. As GAPDH binds to the abundant milk protein lactoferrin (LTF) (Malhotra et al., 2016) and traffics it into cells, this may be a point of interest in future studies exploring changes to the EV surfaceome resulting from heat treatment and lyophilisation.

Overall, these results suggest that the methods used in this study to recover EVs from HM, CM and IF were successful in enriching for EVs, including exosomes. The abundance of EV-enriched proteins seems to be variable between species and sample type (e.g., milk vs IF). Hence, the classes of EVs found in milk may differ between species, or else vesicle populations and/or protein integrity are changed due to IF processing.

### 3.3 Quantitative Proteomic Comparison of HM and CM EVs

To determine whether HM and CM EVs were similar in their protein abundance and complexity, we quantified and compared EV proteins in HM and CM by SWATH-MS. The PCA plot showed clear separation of HM and CM samples, with clustering of two distinct groups away from pooled biological quality control samples (Supplementary file 2, Figure S2). Based on peptides of identical sequence, 216 proteins were quantified and common to HM and CM; 126 were enriched and 17 depleted in CM compared to HM (Figure 2). Among the CM-enriched were immunomodulatory proteins LDH, XDH and BT1A1, G proteins (24% of total) and metabolite interconversion enzymes (22% of total). Thirteen of the 17 proteins more abundant in HM EVs were components of 40S and 60S ribosomal subunits, which have been identified in a separate CM EV study (Benmoussa et al., 2019). LTF was more abundant in HM EVs and although it is an abundant milk protein, it is enriched in EVs via its association with GAPDH (Dar et al., 2021). As GAPDH abundance was similar across HM and CM EV samples, this finding is in agreement with a previous study that found LTF to be 10x more abundant in HM compared to ruminant milk (Lu et al., 2018). However, in this study, LTF in HM EVs was ∼100x more abundant than in CM EVs, representing an overall 10-fold increase in HM EVs compared to LTF found in whole milk. LTF regulates iron homeostasis and contributes to bacterial resistance in the human infant (Ballard & Morrow, 2013), thus such a marked increase of LTF in HM and HM EVs may be of biological significance and warrants further investigation.

**Figure 2:**
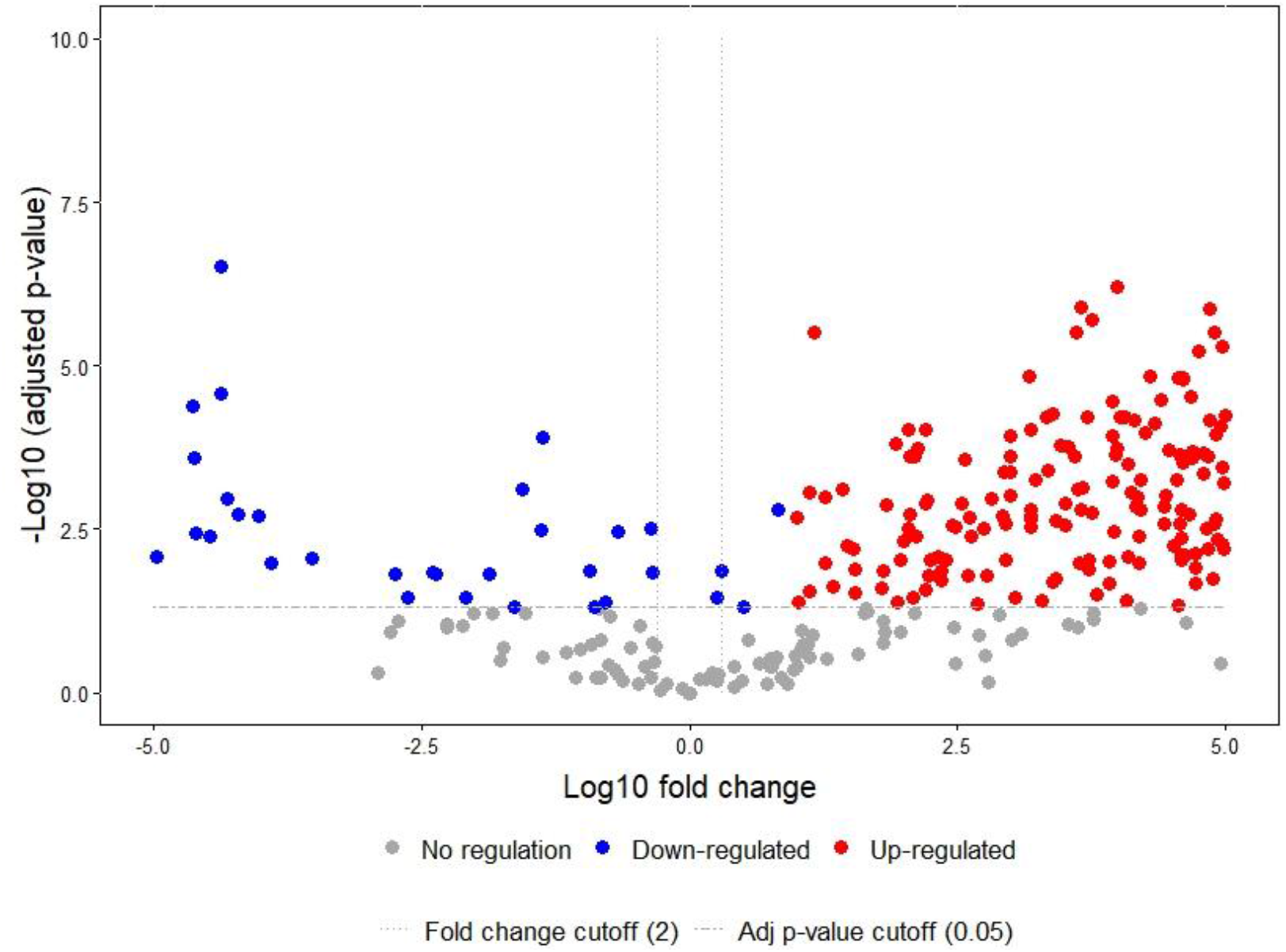
Volcano plot of differential protein abundance in extracellular vesicles (EVs) enriched from human (HM) and cow’s milk (CM). 216 proteins were quantified in total based on peptide sequence homology. Red dots to the right (126) represent proteins more abundant in CM EVs, blue dots to the left (17) represent proteins more abundant in HM EVs.

Rab family proteins were more abundant in CM EVs (11/126), which are associated with EVs arising from multivesicular bodies such as exosomes (Benmoussa et al., 2019). Although Rab proteins are known to play an essential role in directing vesicle transport and targeting vesicles to cell membranes for uptake, their function in milk EVs remains understudied (Blanc & Vidal, 2018).

### 3.4 Qualitative Proteomic Comparison of EVs Recovered from Milk and IF Products

Our workflow utilising SWATH-MS enabled qualitative (identification) analysis of the EV proteome from milk and IF products. EVs isolated from the 9 IF products used in this study resulted in 517 unique proteins, 343 of which were shared by all IF EV samples. Of these 343 proteins, 127 were shared by HM, CM and IF EVs (Figure 3). Out of the proteins that were listed as classified in the PANTHER database, enzymes and binding proteins represented 34% and 44% of the shared proteins by molecular function, respectively (Supplementary file 2, Figure S3). Metabolite interconversion enzymes (17%) and protein-binding activity modulators (23%) were among the most abundant by protein class. As enzymes are related to the breakdown of macromolecules and thus could contribute to digestion of milk proteins, their conservation across species and EV sample types is an important consideration for the infant digestive tract. Differences in the abundance of enzymes or types of enzymes contained in EVs from the different milk types may impact the ability of the infant to optimally digest nutritional components of milk, and thus contributes to optimal growth and development. Indeed, of the proteins *<u>uniquely</u>* identified in HM EVs, proteins with catalytic activity and binding proteins were the most abundant (23% and 39%, respectively), including aminopeptidases which play a known role in the breakdown of proteins via removal of amino acids from the amino end of proteins and peptides (Taylor, 1993). Recent studies have hinted at the need for further macro analysis of HM and IF products due to known compositional differences that could affect enzymatic digestion of IF by the infant gut, including changes to MFG size and fat composition that affects the functional lipid particle surface (Manoni et al., 2020; Yao et al., 2021). In this study, for the first time, we provide evidence that IF EVs contain enzymes, which suggests that encapsulation within EVs may protect enzymes from degradation during IF processing. This information could be leveraged as the EV community moves towards loading biological (i.e., milk-derived) and synthetic lipid nanoparticles with compounds for targeted delivery of molecular cargo (de Abreu et al., 2021).

**Figure 3:**
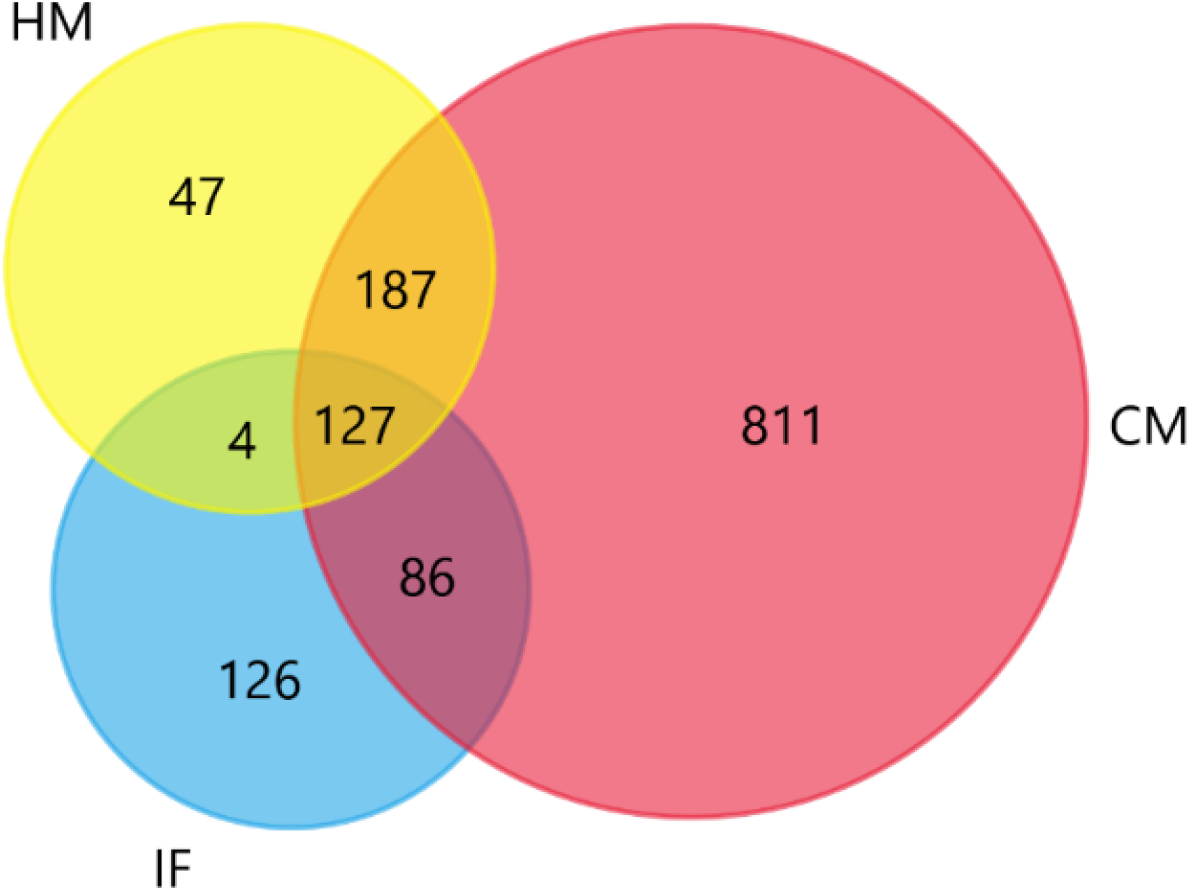
Venn diagram of proteins identified in human milk (HM), cow’s milk (CM) and infant formula (IF) extracellular vesicles.

### 3.5 MiRNAs Recovered from Milk and IF EVs

The total number of miRNAs identified in HM, CM and IF EVs were 832, 1357, and 1489, respectively (Supplementary file 2, Figure S4). Surprisingly, none of the identified miRNA were unique to HM EVs, however 729 were identified in all samples. HM and CM EVs had 103 miRNAs in common that were not identified in IF EVs. These findings are in agreement with a separate study on the similarity of miRNAs found in human, cow and goat milk (Golan-Gerstl et al., 2017). Our data demonstrates that miRNAs found in HM EVs are conserved in CM EVs and a high percentage of miRNAs found in CM EVs are likely to survive IF processing.

Quantitative miRNA analysis of HM and CM EVs found that of the top 50 differentially expressed (DE) miRNA, all 50 were more abundant in HM EVs (Figure 4). Similar trend is evident in the expression of immune related miRNAs between HM and CM EV, miR-148a, miR-181 class (miR-181a-1-3p, miR-181b-2-5p and miR-181a-2-3p) and miR-92b-3p. However, miR-150-3p, a suppressor of B-cell differentiation, was highly expressed in CM EVs compared to HM EVs, which is in agreement with the results of Kosaka and colleagues (2010), who found that HM EVs were not enriched for miR-150-3p (Supplementary file 2, Figure S5). In contrast, the difference between the top 50 DE miRNAs in CM vs IF EVs showed few remarkable differences with most *z*-scores ranging from -1 to 1 (Supplementary file 2, Figure S6), which indicates that milk EVs may preserve miRNAs and act as a protective barrier during IF production. Interestingly, however, there was no overlap between the top 50 DE miRNA from HM vs CM and CM vs IF EVs. These findings suggest that HM EVs may be enriched with miRNA related to immune differentiation and development that differs from that found in CM EVs, which could be of interest in future studies on adapting IF products for susceptible infant groups, such as pre-term infants.

**Figure 4:**
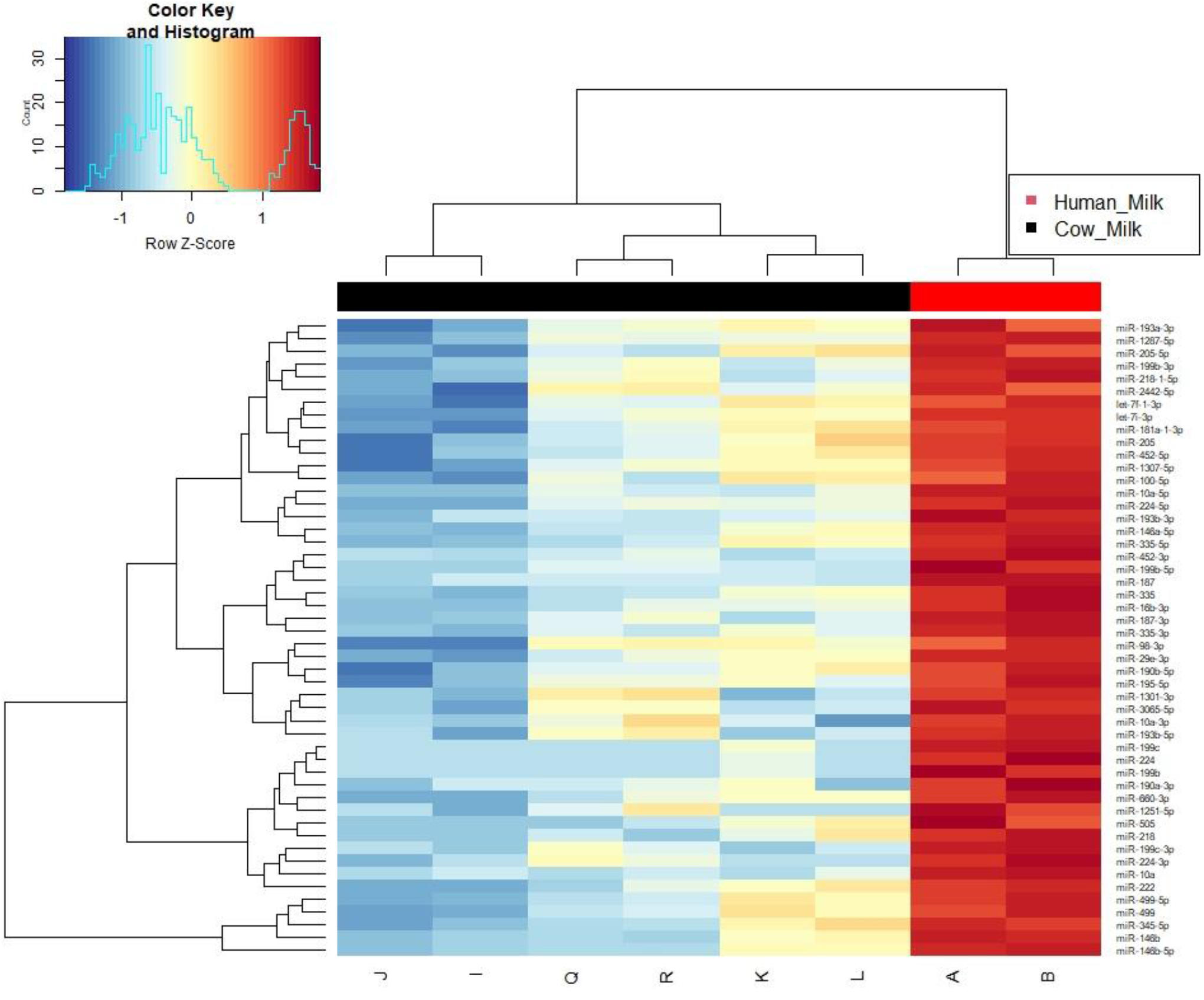
Heatmap of the top 50 differentially expressed (DE) microRNAs found in extracellular vesicles (EVs) enriched from human milk (HM) and cow’s milk (CM). Columns A and B = HM EVs; columns I – L, Q and R = CM EVs.

Using miRNA target prediction tools, the top 50 DE miRNA in HM vs CM EVs were mapped to 9107 gene targets (Supplementary file 6). Gene ontology analysis of these gene targets revealed association with the following protein classes; protein modifying enzymes (13%), metabolite interconversion enzymes (14%), and gene-specific transcriptional regulators (13%) (Supplementary file 2, Table S2). Using the same target prediction tools, 712 gene targets were identified from the top 50 DE miRNA between CM and IF EVs (Supplementary file 7). Although the top 50 DE miRNAs from HM vs CM and CM vs IF EVs did not overlap, there were 436 gene targets common to DE miRNAs from both comparisons (Supplementary file 2, Figure S7). GO analysis by molecular function of these gene targets found that of the classified genes, 32% were associated with catalytic activity and 39% with binding (Supplementary file 2, Table S3). These findings correlate with the proteomics data in the current study, which highlights a potential link between the EV transcriptome and proteome of milk and IF samples. Milk EV-derived miRNAs may therefore be driving transcriptional regulation of enzymes associated with a wide variety of cellular functions, albeit via different miRNAs dependent upon their species of origin, and possibly via different mechanisms.

MiRNAs found in milk have been described as resistant to the harsh conditions of the digestive tract, owing to their encapsulation within EVs (Benmoussa et al., 2020). A study by Liao and colleagues (2017) modelled gastric digestion of milk EVs *in vitro* and found that digested and undigested EVs enter human intestinal epithelial crypt-like cells, with approximately 10% of milk EVs localising to the cell nucleus (Liao et al., 2017). Differential abundance of EV miRNA between HM and CM may therefore provide critical information regarding genetic regulation during what is arguably the most developmentally susceptible period of human life.

### 3.6 Conclusions

This study is the first to apply an established method of milk EV isolation for the enrichment and characterisation of lyophilised EVs in IF products. The successful recovery of IF EVs led to the development of a workflow that would allow cross-species comparison of proteins and miRNA found in HM, CM and IF EVs. The application of this workflow enabled the identification of key species-specific differences in proteins and miRNA quantified in HM and CM EVs and established a link to EVs found in IF products. These differences may assist in optimising nutritional products for formula-fed infants and can be the subject of further investigation.

### CRediT authorship contribution statement

**Natalie P. Turner:** Investigation, Data curation, Formal analysis, Writing – original draft, Writing – Review & editing, Visualisation. **Pevindu Abeysinghe:** Formal analysis, Data curation, Investigation, Writing – Review & editing, Visualisation. **Pawel Sadowski:** Supervision, Resources, Writing – Review & editing. **Murray D. Mitchell:** Conceptualization, Methodology, Resources, Supervision, Writing – Review & editing, Project administration, Funding Acquisition.

## Acknowledgements

This work was supported by Reckitt Benckiser/Mead Johnson (2019-2022). N.P.T. and P.A. are supported by a student scholarship from the Australian Research Council grant (ARC LP160101854) and QUT HDR Tuition fee sponsorship. This work was enabled by use of the Central Analytical Research Facility (CARF) at the Queensland University of Technology (QUT).

## Data availability

We have submitted all relevant data of our experiments to the EV-TRACK knowledgebase (EV-TRACK ID: EV230025) (Van Deun et al., 2017).

The miRNA dataset associated with this work has been submitted to the Vesiclepedia online compendium (Kalra et al., 2012) (http://microvesicles.org/, accessed on 6^th^ February 2023).

All DIA raw files (wiff/.wiff.scan) DIA-NN output files (.xlsx) have been deposited to the ProteomeXchange Consortium via the PRIDE (Collins et al., 2017) partner repository (http://www.proteomexchange.org/, accessed on 7^th^ February 2023) with the dataset identifier PXD039923.

## Abbreviations

ALB: Bovine serum albumin
AMBIC: Ammonium bicarbonate
BTN1A1: Butyrophilin subfamily 1 member A1
CALNX: Calnexin
CM: Cow’s milk
DDA: Data-dependent acquisition
DIA: Data-independent acquisition
DP: Declustering potential
DPBS: Dulbecco’s phosphate buffered saline
EV: Extracellular vesicle
FDR: False discovery rate
FLOT-1: Flotillin-1
GAPDH: Glyceraldehyde-3-phosphate dehydrogenase
GO: Gene ontology
HM: Human milk
IF: Infant formula
iRT: indexed retention time
JEG: JEG-3 choriocarcinoma cell lysate
LC: Liquid chromatography
LDH: Lactadherin
LTF: Lactoferrin
MFG: Milk fat globule
miRNA: microRNA
MS: Mass spectrometry
OREI: Office of Research Ethics & Integrity
PBQC: Pooled biological quality control
PBS: Phosphate buffered saline
PBST: Phosphate buffered saline/0.1% Tween-20
PCA: Principal component analysis
PLIN2: Perilipin-2
SCX: Strong cation exchange
SEC: Size-exclusion chromatography
SWATH-MS: Sequential Window Acquisition of all Theoretical Mass Spectra
SYN-1: Syntenin-1
TEM: Transmission electron microscopy
TFA: Trifluoroacetic acid
TOF: Time-of-flight
TSG101: Tumour susceptibility gene 101
UC: Ultracentrifugation
WB: Western blot
XDH: Xanthine dehydrogenase/oxidase

