## Supplementary file 2 for "Cross-species proteomic and microRNA comparison of extracellular vesicles in human milk, cow’s milk, and infant formula products: moving towards next generation infant formula products"

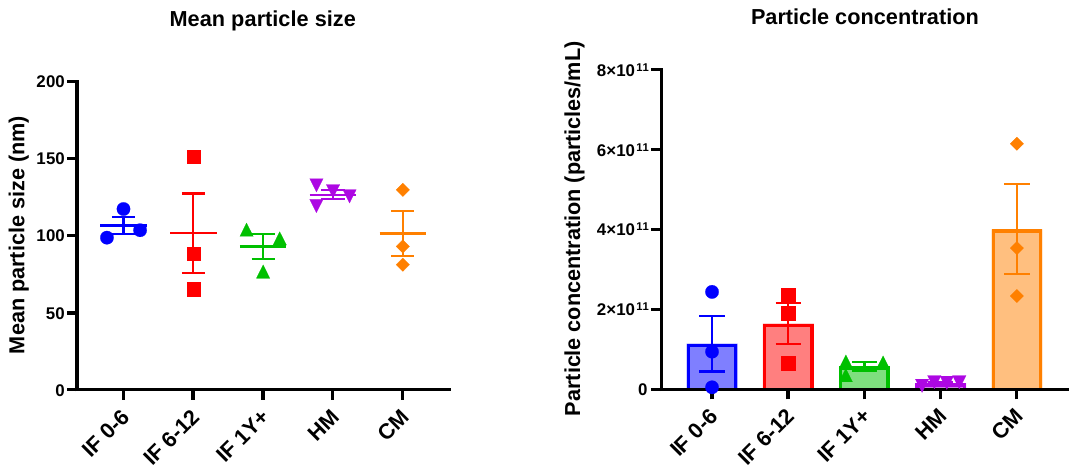

**A**

**B**

**Figure S1:** A: Mean particles size in extracellular vesicle (EV) samples enriched from infant formula (IF) (*n* = 3/group), human milk (HM) (*n* = 4) and cow’s milk (CM) (*n* = 3). B: Particle concentration of EV samples from IF, HM and CM. 0-6 = 0–6-month formula; 6-12 = 6-12-month formula; 1Y+ = 1 year and over formula. Error bars = mean ± SEM.

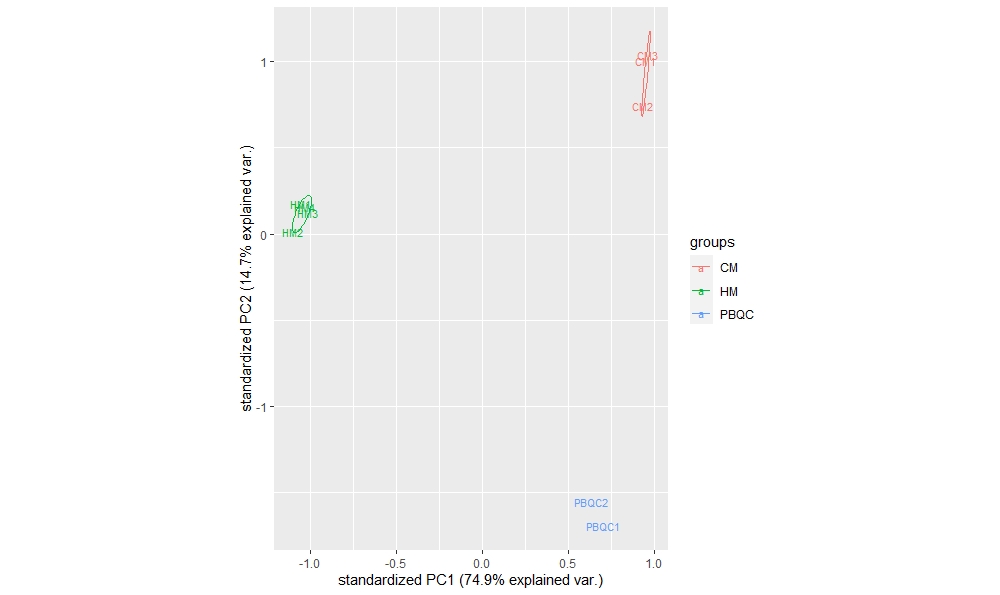

**Figure S2:** Principal component analysis plot of quantifiable proteins identified in extracellular vesicles enriched from human milk (HM), cow’s milk (CM) and a pooled biological quality control sample (PBQC).

**Figure S3:** Proteins common to extracellular vesicles enriched from human milk (HM), cow’s milk (CM) and infant formula (IF) products by protein class.

|  |  |  |  |
| --- | --- | --- | --- |
| 1 transporter activity (GO:0005215) | 6 | 4.7% | 5.7% |
| 2 translation regulator activity (GO:0045182) | 1 | 0.8% | 0.9% |
| 3 catalytic activity (GO:0003824) | 36 | 28.3% | 34% |
| 4 molecular function regulator (GO:0098772) | 3 | 2.4% | 2.8% |
| 5 ATP-dependent activity (GO:0140657) | 1 | 0.8% | 0.9% |
| 6 low-density lipoprotein particle receptor activity (GO:0005041) | 1 | 0.8% | 0.9% |
| 7 molecular transducer activity (GO:0060089) | 3 | 2.4% | 2.8% |
| 8 molecular adaptor activity (GO:0060090) | 6 | 4.7% | 5.7% |
| 9 structural molecule activity (GO:0005198) | 2 | 1.6% | 1.9% |
| 10 binding (GO:0005488) | 47 | 37% | 44.3% |

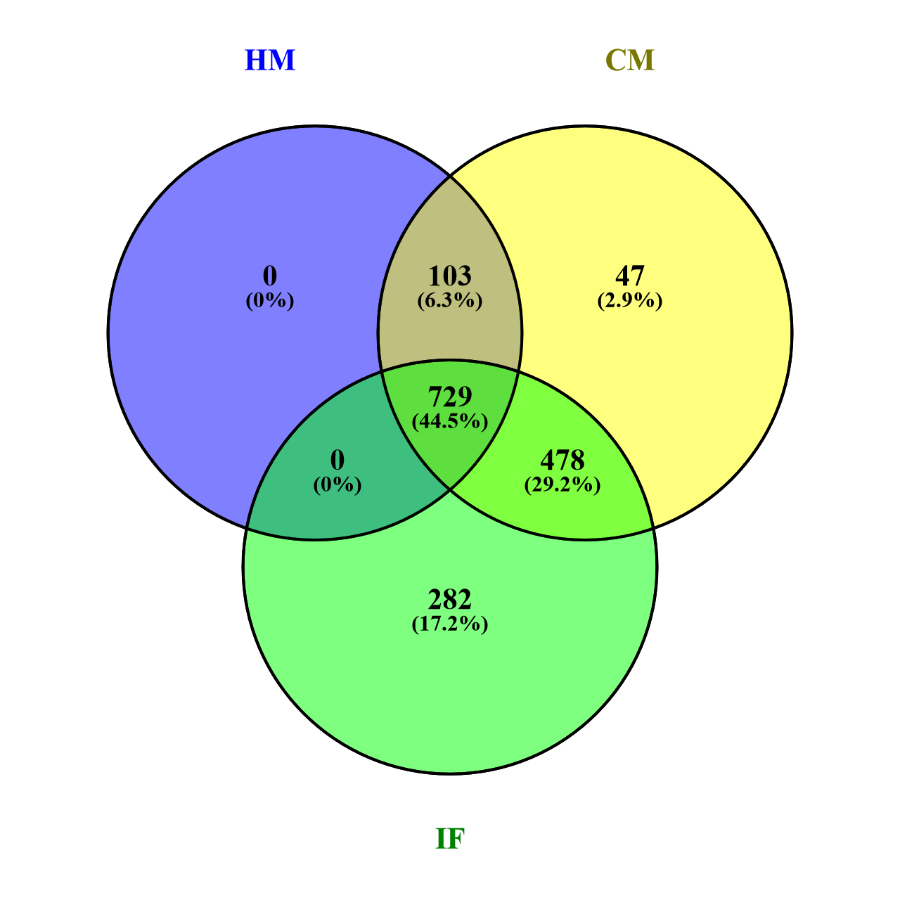

**Figure S4:** Venn diagram of micro(mi)RNAs identified in extracellular vesicles enriched from human milk (HM), cow’s milk (CM) and infant formula (IF) products.

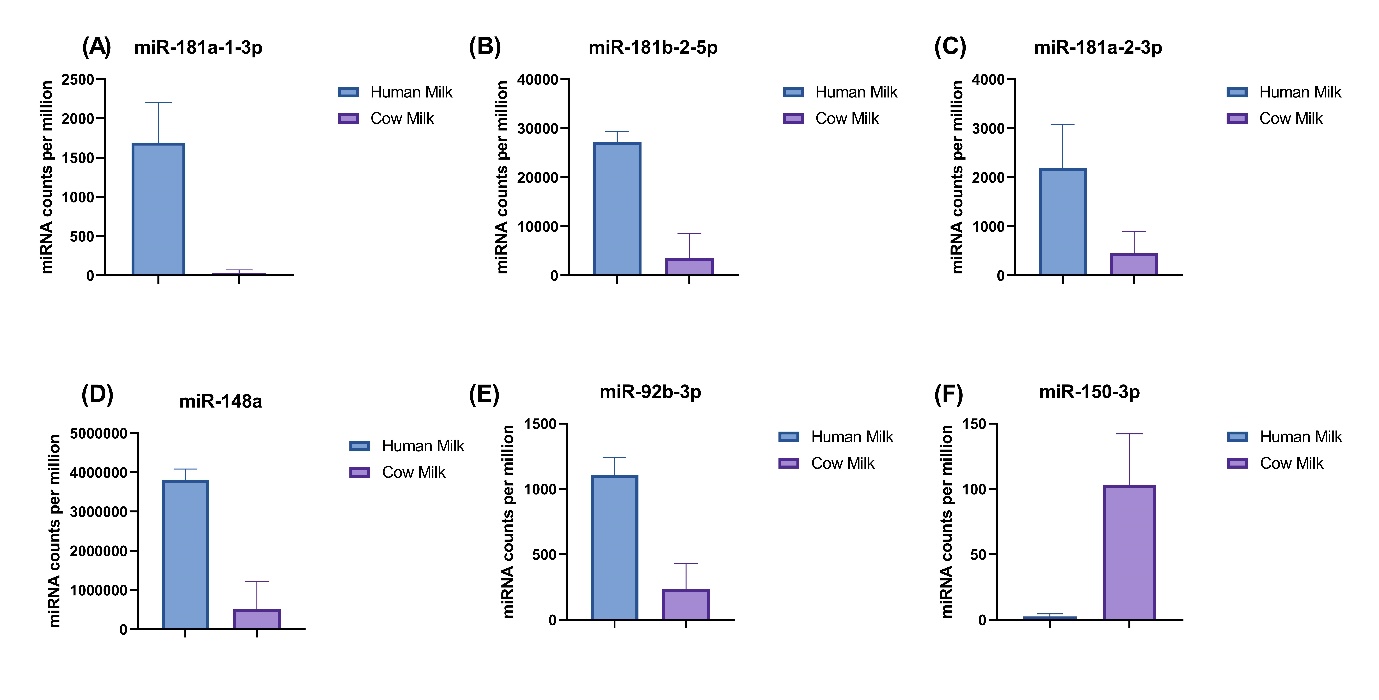

**Figure S5:** Immune related micro(mi)RNA expression levels between human (HM) and cow milk (CM) EVs. HM contains higher miRNA counts in miR-181 class (A, B and C), miR-148a (D) and miR-92b-3p (E). However, miR-150-3p was highly expressed in CM EVs.

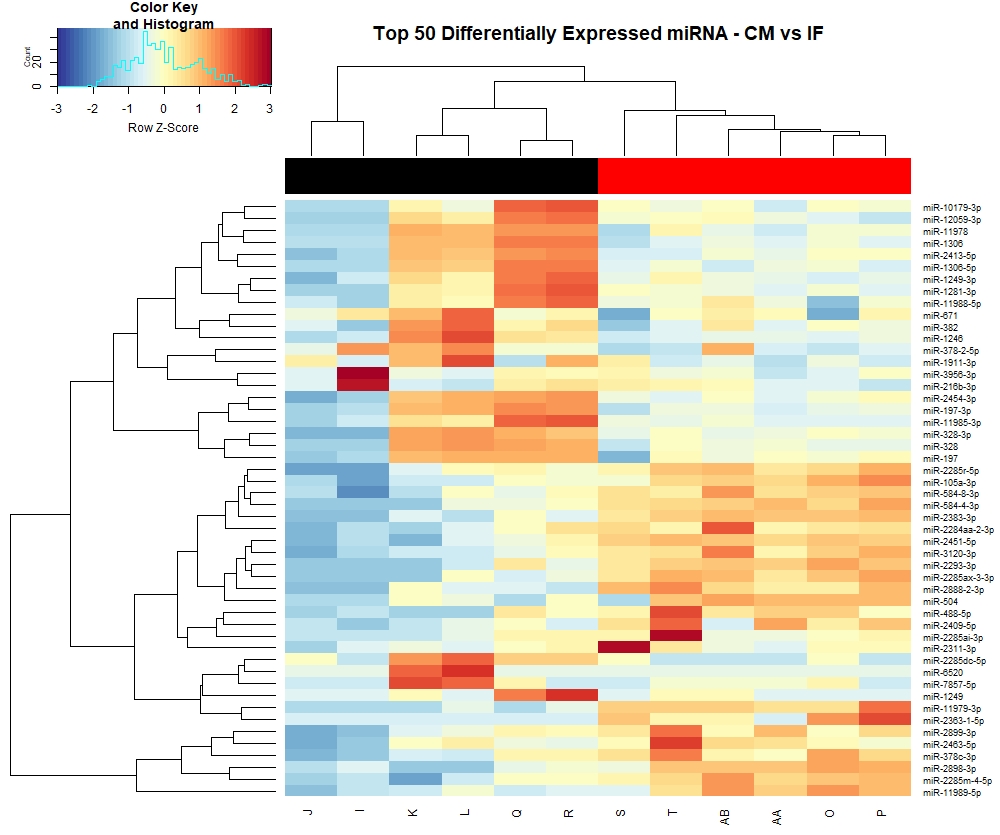
**Figure S6: Heatmap of the top 50 differentially expressed micro(mi)RNA identified in extracellular vesicles recovered from cow’s milk (CM) and infant formula (IF) products.** Columns J, I, K, L, Q and R = CM; columns S, T, AA, AB, O and P = IF.

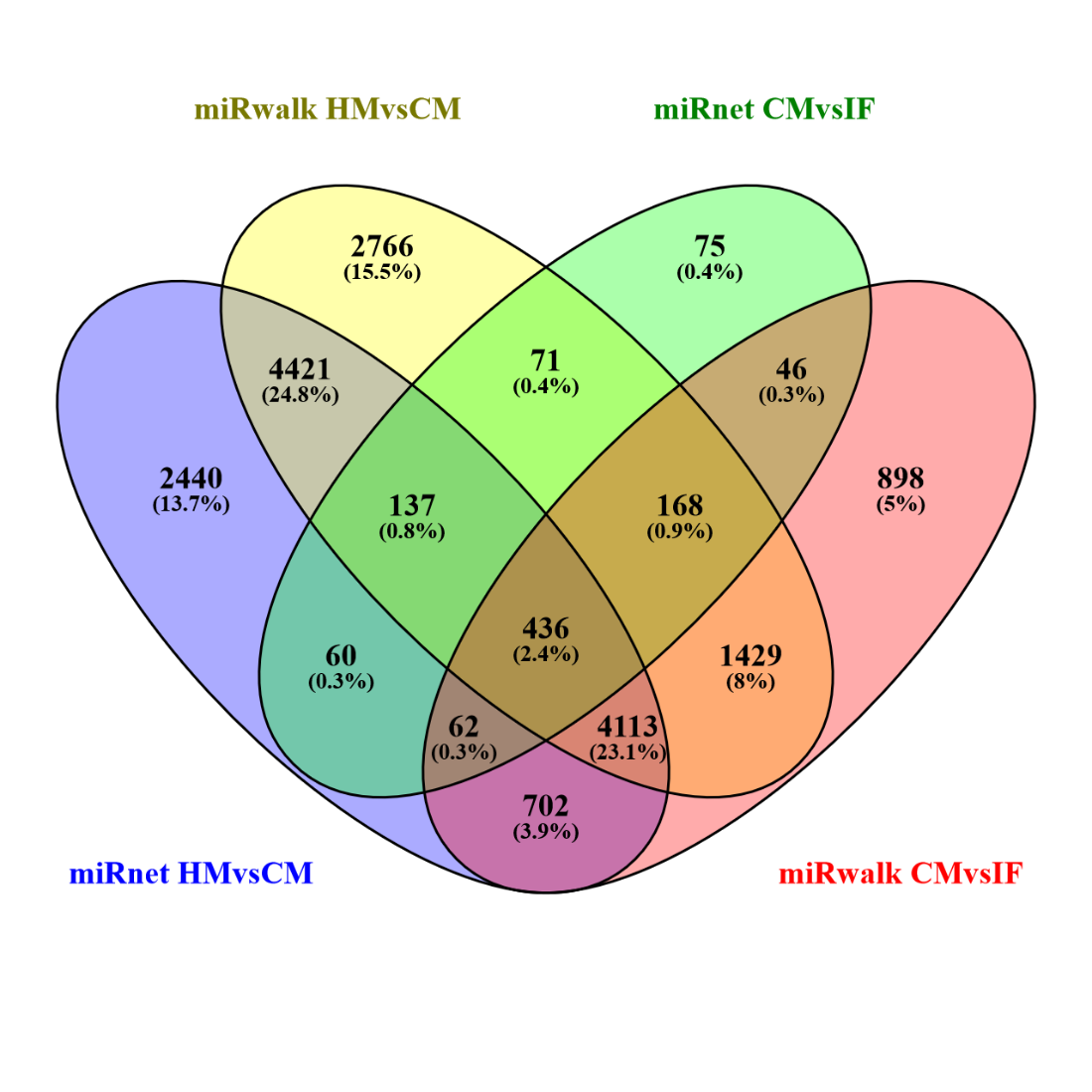

**Figure S7:** Venn diagram of the predicted micro(mi)RNA targets linked to the top 50 differentially expressed miRNA identified in extracellular vesicles from human milk (HM) cow’s milk (CM) and infant formula (IF) products.

**Table S1: Particle and protein yields for Human milk (HM), cow’s milk (CM) and infant formula (IF) samples obtained from nanoparticle tracking analysis.** Values are based on the total volume of the pooled size-exclusion chromatography fractions (4 x 500 µL fractions = 2 mL). *Average of technical replicates given.

| **Sample** | **Particle yield** | **Protein Yield (µg)** | **Particles/µg protein** | **Particles/mL starting volume** |
| --- | --- | --- | --- | --- |
| HM* | 3.09E+10 | 114 | 2.71E+08 | 1.72E+08 |
| CM* | 1.23E+12 | 547 | 2.25E+09 | 6.15E+09 |
| IF1-0-6-month | 1.89E+11 | 32000 | 5.91E+06 | 5.25E+08 |
| IF2-0-6-month | 1.05E+10 | 3338 | 3.15E+06 | 5.25E+07 |
| IF3-0-6-month | 4.88E+11 | 3205 | 1.52E+08 | 2.44E+09 |
| IF1-6-12-month | 1.33E+11 | 32000 | 4.16E+06 | 3.69E+08 |
| IF2-6-12-month | 3.80E+11 | 3434 | 1.11E+08 | 1.90E+09 |
| IF3-6-12-month | 4.74E+11 | 3057 | 1.55E+08 | 2.37E+09 |
| IF1-1 year + | 1.35E+11 | 31236 | 4.32E+06 | 3.75E+08 |
| IF2-1 year + | 1.40E+11 | 2895 | 4.84E+07 | 7.00E+08 |
| IF3-1 year + | 7.72E+10 | 346 | 2.23E+08 | 3.86E+08 |

**Table S2: Gene ontology (GO) results by protein class of miRNA gene targets common to Human and cow’s milk EVs.**

| **#** | **GO Accession: Protein Class** | **# Of genes** | **% Total** |
| --- | --- | --- | --- |
| 1 | extracellular matrix protein (PC00102) | 57 | 0.90% |
| 2 | cytoskeletal protein (PC00085) | 283 | 4.30% |
| 3 | transporter (PC00227) | 534 | 8.10% |
| 4 | scaffold/adaptor protein (PC00226) | 398 | 6.10% |
| 5 | DNA metabolism protein (PC00009) | 133 | 2.00% |
| 6 | cell adhesion molecule (PC00069) | 152 | 2.30% |
| 7 | intercellular signal molecule (PC00207) | 169 | 2.60% |
| 8 | protein-binding activity modulator (PC00095) | 450 | 6.90% |
| 9 | viral or transposable element protein (PC00237) | 10 | 0.20% |
| 10 | RNA metabolism protein (PC00031) | 431 | 6.60% |
| 11 | calcium-binding protein (PC00060) | 51 | 0.80% |
| 12 | gene-specific transcriptional regulator (PC00264) | 848 | 12.90% |
| 13 | defense/immunity protein (PC00090) | 123 | 1.90% |
| 14 | translational protein (PC00263) | 150 | 2.30% |
| 15 | metabolite interconversion enzyme (PC00262) | 935 | 14.30% |
| 16 | protein modifying enzyme (PC00260) | 864 | 13.20% |
| 17 | chromatin/chromatin-binding, or -regulatory protein (PC00077) | 163 | 2.50% |
| 18 | transfer/carrier protein (PC00219) | 56 | 0.90% |
| 19 | membrane traffic protein (PC00150) | 244 | 3.70% |
| 20 | chaperone (PC00072) | 97 | 1.50% |
| 21 | cell junction protein (PC00070) | 25 | 0.40% |
| 22 | structural protein (PC00211) | 43 | 0.70% |
| 23 | storage protein (PC00210) | 2 | 0.00% |
| 24 | transmembrane signal receptor (PC00197) | 339 | 5.20% |

**Table S3: Gene ontology (GO) results by molecular function of miRNA gene targets common to Human, cow’s milk, and infant formula EVs.**

| **#** | **GO Accession: Molecular Function** | **# Of genes** | **% Total** |
| --- | --- | --- | --- |
| 1 | transporter activity (GO:0005215) | 28 | 7.70% |
| 2 | translation regulator activity (GO:0045182) | 5 | 1.40% |
| 3 | transcription regulator activity (GO:0140110) | 26 | 7.10% |
| 4 | catalytic activity (GO:0003824) | 115 | 31.50% |
| 5 | cytoskeletal motor activity (GO:0003774) | 2 | 0.50% |
| 6 | molecular function regulator (GO:0098772) | 19 | 5.20% |
| 7 | ATP-dependent activity (GO:0140657) | 5 | 1.40% |
| 8 | molecular transducer activity (GO:0060089) | 17 | 4.70% |
| 9 | molecular adaptor activity (GO:0060090) | 3 | 0.80% |
| 10 | structural molecule activity (GO:0005198) | 3 | 0.80% |
| 11 | binding (GO:0005488) | 142 | 38.90% |
