## Supplementary file 3 for "Cross-species proteomic and microRNA comparison of extracellular vesicles in human milk, cow’s milk, and infant formula products: moving towards next generation infant formula products"

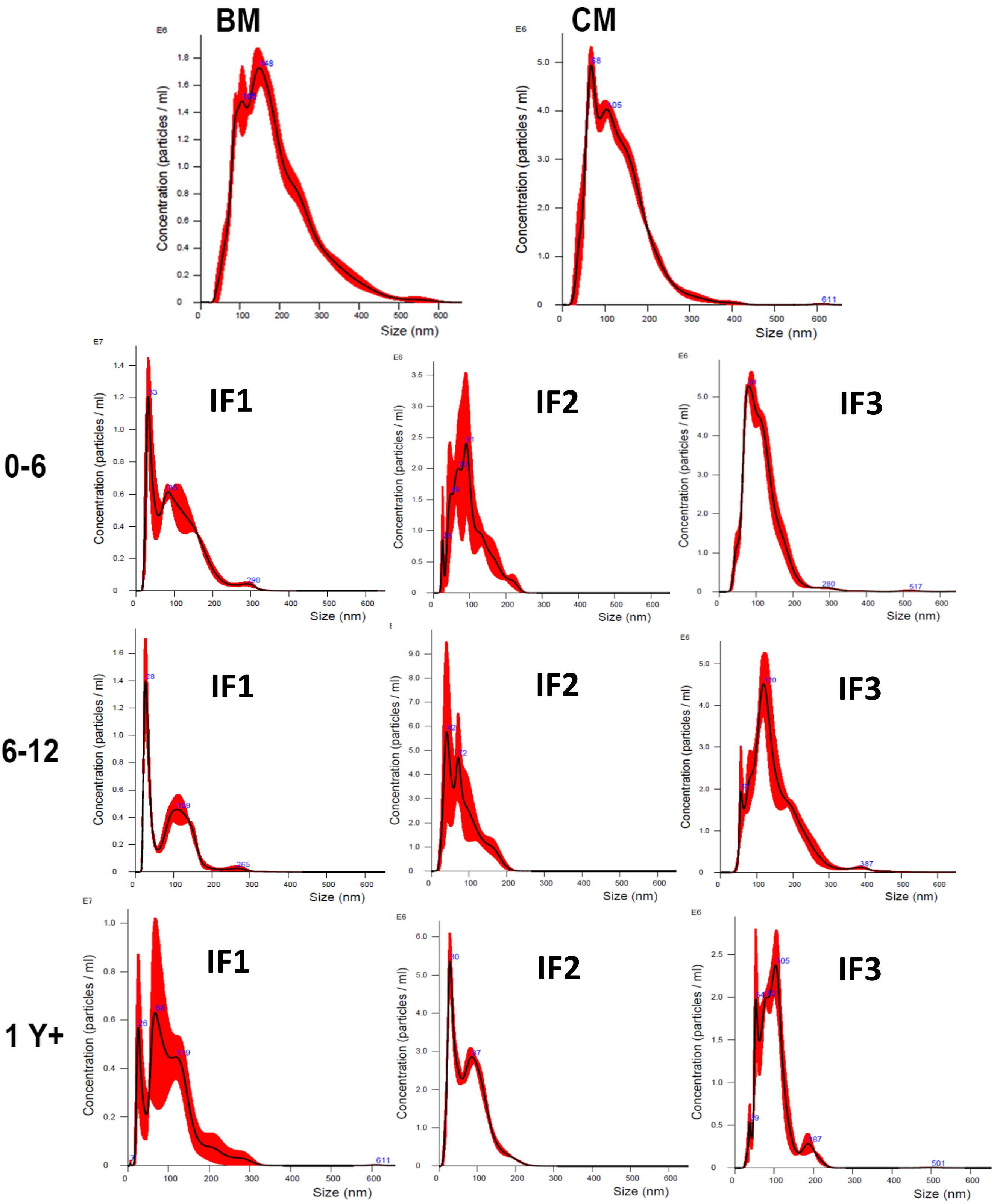


**Nanoparticle tracking analysis size distribution plots for Human milk (HM), cow's milk (CM) and infant formula (IF) products.** X axis = particle size (nm); Y axis = particle concentration (particles/mL x$10^{6}$). Data presented are raw values (i.e., not multiplied by dilution factors).
