## Supplementary file 4 for "Cross-species proteomic and microRNA comparison of extracellular vesicles in human milk, cow’s milk, and infant formula products: moving towards next generation infant formula products"

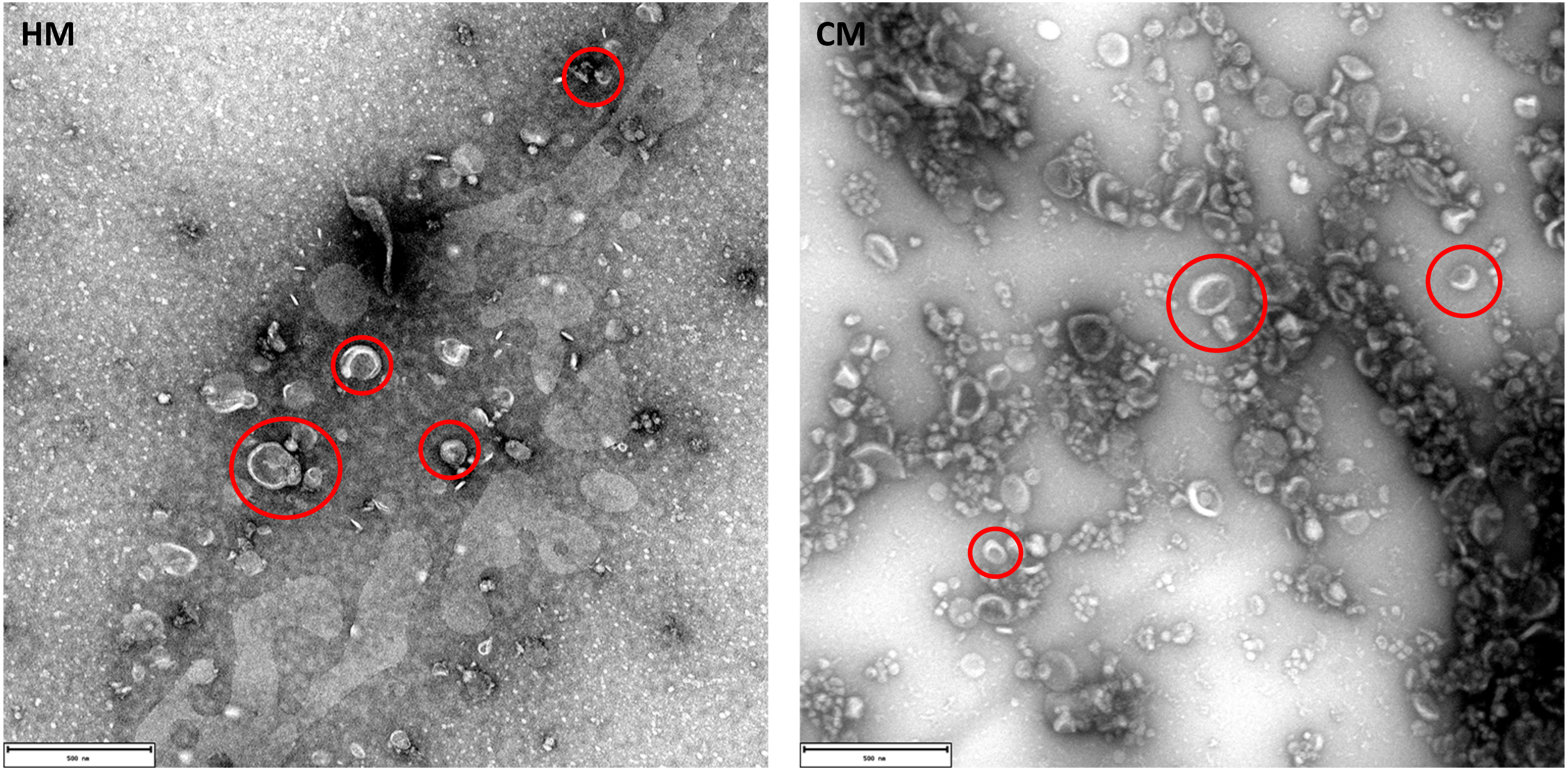


**Figure S1: Transmission electron microscopy representative images of human milk (HM; left panel) and cow's milk (CM; right panel) extracellular vesicles (pooled fractions 7 – 10).** Scale bar = 500 nm. Red circles indicate vesicles of various sizes (range ~30 – 200 nm).

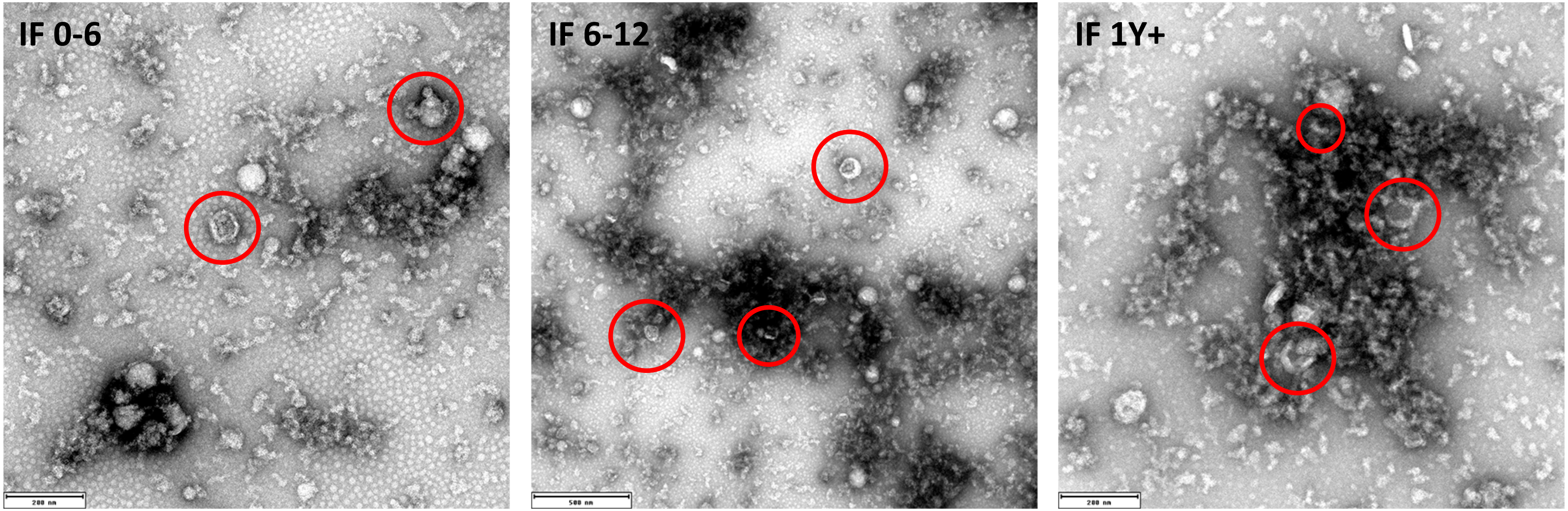


**Figure S2: Transmission electron microscopy representative images of extracellular vesicles (pooled fractions 7 – 10) recovered from 0-6-month infant formula (IF; left panel), 6-12-month IF (middle panel), and 1 year and over IF (right panel).** Scale bar (left and right panels) = 200 nm; scale bar (middle panel) = 500 nm. Red circles indicate vesicles of various sizes (range ~50 – 200 nm).
